# Modeling and analysis of site-specific mutations in cancer identifies known plus putative novel hotspots and bias due to contextual sequences

**DOI:** 10.1101/2020.02.07.939512

**Authors:** Victor Trevino

## Abstract

In cancer, recurrently mutated sites in DNA and proteins, called *hotspots*, are thought to be raised by positive selection and therefore important due to its potential functional impact. Although recent evidence for APOBEC enzymatic activity have shown that specific types of sequences are likely to be false, the identification of putative hotspots is important to confirm either its functional role or its mechanistic bias. In this work, an algorithm and a statistical model is presented to detect hotspots. The model consists of a *beta-binomial* component plus fixed effects that efficiently fits the distribution of mutated sites. The algorithm employs an optimal step-wise approach to find the model parameters. Simulations show that the proposed algorithmic model is highly accurate for common hotspots. The approach has been applied to TCGA mutational data from 33 cancer types. The results show that well-known cancer hotspots are easily detected. Besides, novel hotspots are also detected. An analysis of the sequence context of detected hotspots show a preference for TCG sites that may be related to APOBEC or other unknown mechanistic biases. The detected hotspots are available online in http://bioinformatica.mty.itesm.mx/HotSpotsAnnotations.

## Introduction

It is thought that recurrently mutated amino-acid positions in cancer genes, namely mutation hotspots, are likely to have an important functional impact [1]. Several well-known examples support this view. One of the most frequent hotspots, BRAF V600E mutation, is known to overactivate the RAS pathway [2,3]. BRAF is top mutated in thyroid carcinoma [4], melanoma [5], and hairy-cell leukemia [3], and also frequent in colon and lung cancers [6–8]. Other hotspots are also well-known such R132H in IDH1 for low-grade gliomas [9], G12/G13 in KRAS for lung [10], and Q61 in NRAS for melanoma [11]. Many other genes also show hotspots [12].

Some non-cancer genes seem to show hotspots that become clear when mutations from all cancers are aggregated [1,12,13]. For example, in Chang *et al.* analysis [13], the RRAS2 showed a hotspot in Q72, which is still not marked as a cancer gene in the Cosmic curated database revision 2019 [14] neither detected for positive selection in Martincorena analysis [15]. This suggests that the identification of putative novel hotspots is important in cancer.

Some methods have been reported regarding the detection of mutation hotspots. Of the seminal approaches, there was a tendency to identify regions [16,17] or domains [1,18] when the available mutations were more limited. Similarly, some approaches focused on the threedimensional protein structure to identify mutation-rich 3D-regions [19–21]. Then, position-specific models were proposed [12,13,22,23]. These approaches used a binomial or a Poisson distribution to model mutation distribution across genes. Nevertheless, the mutation distribution per gene may depend on cofactors such as sequence context [12], gene length [24], cancer type [25,26], mutational processes [26,27], or relative position along nucleosomes [28]. Modeling all these cofactors together is a very difficult task given its complexity and lack of data to sufficiently estimate embed parameters. To account for these and other unknown factors, an over-dispersion model is preferred [15,24,29]. Thus, other approaches utilize more appropriate models such as the beta-binomial model [24,29], which were applied to non-coding regions.

Although the above methods have been useful, there are some pitfalls. Some approaches use binomial or Poisson models with one or two cofactors [12,13,22] but this may lead to many false positives or negatives. For example, there are 20 genes reported by Chang *et al.* [13] that show significant hotspot associations backed up by only two mutations such as SESN2. The certainty to pursue experimental validations on SENS2 is very difficult if there is no available information about the parameters fitted and all related information used regarding the gene and mutations. Some methods use randomization of the mutations to estimate significance [1] but this would lead to biased estimations if not all cofactors are considered, which is difficult because there is still uncertainty about possible cofactors. Other methods use well-known cancer genes as positive controls and presumed negatives to estimate sensitivity and specificity [30]. One of the reported problems of this strategy is that it sacrifices sensitivity for specificity [30], which may show difficulties when used as a discovery tool. In this context, simulations may be a good strategy.

One of the strengths of methods that detect regions, domains, and 3D structures is that estimations can be more reliable because many more mutations can be analyzed within regions than within positions. Nevertheless, this is also a weakness because it is known that the sequence context plays a role [12] and these methods lack nucleotide sequence resolution. Another issue is that some methods focus on single nucleotide variants, presumably because of the lack of corrections for small insertions and deletions (INDELS) [13]. Regarding the types of mutations, most referred methods focus mainly on missense mutations. This is sensible because these hotspots of this mutation type mark positions on the protein that may change its function. Besides, missense mutations represent a large proportion of all mutations. Nevertheless, other mutations may be interesting such as those generated by small insertions and deletions that may easily accumulate at repetitive sequences [31]. A deeper analysis of methods is presented elsewhere [32].

In this work, firstly, a comparison of the fitting of the distribution of all types of small mutations by two canonical distributions (*Binomial, Geometric*) and two more that consider overdispersion (*Beta-Binomial,* and *Zero-Inflated Beta-Binomial*) is presented. The comparison leads to the determination that, overall, the *beta-binomial* model seems to be the best model. Then, to account for genuine hotspots that do not fit well even considering over-dispersion by the b*eta-binomial*, a mixture model with fixed effects is proposed to better fit the observed mutation distribution per gene without covariates. The need of fixed effects on high frequent mutations suggests the presence of hotspots. Simulations show that the proposed mixture model is accurate. Then, the mixture model has been applied to *The Cancer Genome Atlas* (TCGA) dataset and the putative hotspots are analyzed. The analysis shows that there is a bias for a sequence context centered at the mutation position and that systematic bias is observed in most co-localized olfactory receptors and other co-localized gene families. More importantly, some detected genes not considered as mutations hotspots show comparable statistics that current well-known cancer genes carrying hotspots. To the author knowledge, this is one of the few methods that use simulations to evaluate the sensitivity and specificity of the proposed method.

## Material and Methods

### Mutational data

The mutation annotation files (maf) were obtained from the public cancer repository TCGA (http://firebrowse.org/) in January 2018 corresponding to 33 cancer types, 10,182 patients, and 3,175,929 mutations (Supplementary Table 1). Only mutations annotated to an amino acid position within its corresponding transcript were used.

### Distribution of mutated positions

For each gene, the mutations were counted per amino acid position depending on their corresponding transcript and protein. Then, the number of amino acid positions having *m_g,i_* mutations (from *0* to *M_g_*) were aggregated where *g* is the gene, *i* is the number of mutations, and *M_g_* is the maximal number of mutations of gene *g* at any amino acid position.

### Distribution Models

To find the optimal parameters to fit a distribution model to the histogram of mutational data, a numerical method implemented in the *optim* function from the *stat* package was used minimizing the difference to the observed distribution (method=“L-BFGS-B” for function *optim* in *stats* package in R, https://cran.r-project.org/). To estimate the difference between fitted and observed distribution was based on the *G-test* statistic, *G* = 2Σ *o_i_* log(*o_i_*/*e_i_*), which is equivalent to the Kullback–Leibler divergency metric used to compare distributions. The geometric and binomial distributions were fitted using the *stat* package in R. The Zero-inflated beta-binomial (ZIBB) was fitted using the *gamlss* package in R. The beta-binomial was fitted using the *emdbook* package in R.

### Beta-binomial model with fixed effects

Conceptually, the problem is schematized in Figure 1A while the algorithm is shown in Figure 1B. The model, *M* = *BetaBin*(*α, β)* +*F*, assumes a fixed effect on positions with an excess of mutations presumably due to hotspots where *M_k_* is the number of positions carrying *k* mutations, *F* is the fixed hotspot effect vector, and *BetaBin* is the beta-binomial density function scaled conveniently to sum the total number of mutations minus the sum of *F*. A stepwise algorithm was devised to fit this model. The algorithm starts setting *F_k_*=0 and fitting the *beta-binomial* model using an optimization algorithm as described in previous section. Then, a matrix of improvements is estimated where each cell represents an independent possible fixed effect in a mutation number *k* (in columns) and at a fraction of the total number of sites (in rows). The value of the cell is the ratio of the *G* statistic before applying the fixed effect divided by the *G* statistic after applying the representing fixed effect. The largest ratio represents an improvement if larger than 1 and therefore it is taken. The corresponding level (positions) and number of mutations *f_i,k_* are aggregated to the *F* vector of fixed effects. The algorithm continues until the largest ratio is not greater than 1 (no improvement), when the number of steps is larger than 3 times the maximum number of mutations, or when the *G* statistic is lower than 1 to avoid over-fitting. The 0 positions (*k*=0), the zero mutations (*m_g,i_*=0), or fractions that do not achieve at least 1 mutation in any *k* mutations, are not explored. The output of the algorithm is the fixed effect vector *F* representing the mutations and the magnitude (number of positions), *F_k_*, that cannot be explained by the *beta-binomial* model with updated *α*, and *β* parameters. The algorithm was implemented in R and is available upon request.

**Figure 1.**
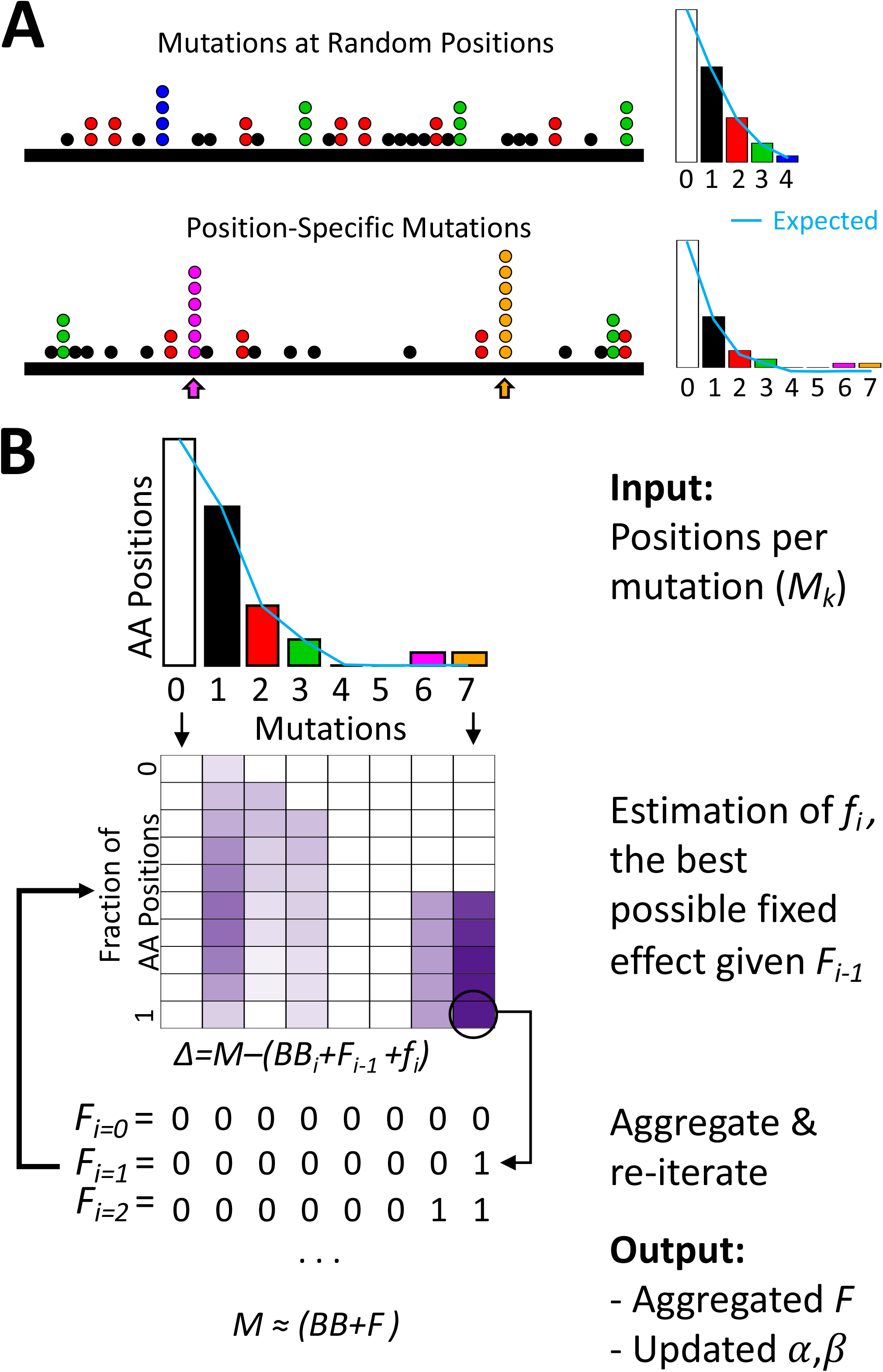
Hotspot concept and proposed algorithm. Panel A Cartoon conceptualization of random mutations along a protein (top) and similar number of mutations resulting in two hotspots (bottom). Histogram of sites per number of mutations is shown. Count are missing for clarity. Panel B shown the proposed algorithm to find the optimal parameters of the mixed model that better fit the observed mutation distribution.

### Simulations

For simulations, sampling with the same protein size and mutations from the beta-binomial model with parameters a and b was used. Then the *F* vector was added depending on the simulation. For no hotspots, *F_k_*=0, otherwise some *F_k_* > 0. In any case, after running the proposed algorithm, a hotspot was recognized if the fitted value of *F_k_* is larger than 50% of the mutations at *k*. From the 2,000 genes taken for simulations, only 1,973 genes generated successful distributions.

### Hotspots from cancer data

For cancer data, a hotspot or biased position was recognized if the fitted value of *F_k_* is larger than 50% of the mutations, whose mutations were 4 or more, and whose q-value (corrected p-value) was <= 0.01. These criteria were used to avoid calling hotspots in low positions that helped to improve model fitting but unlikely to represent hotspots (see Supplementary Figure 1).

### Sequence context

The context sequence of a mutation was annotated using the R package *BSgenome.Hsapiens.NCBI.GRCh38*.

## Results

### Comparisons of competing distributions

To determine the best canonical distribution matching the observed mutations distributions in cancer, a comparison was performed between binomial, geometric, beta-binomial, and zero-inflated beta-binomial (ZIBB) [33]. For this, the Kullback-Leiber divergency metric was used to determine which distribution provides the best fit to the observed distribution. The ZIBB was included due to the observation that sites at zero mutations seem to be exacerbated. Under randomness, the binomial is the expected result. Nevertheless, the results show that the *betabinomial* and the *geometric* functions capture the largest number of genes (Supplementary Figure 2A). The former is expected because the beta-binomial can capture over-dispersion commonly present in binomial data [34]. However, the *geometric* distribution performed surprisingly high. Then, to assess whether there is a preference of a density function for cancer genes, the same process was performed for cancer genes according to Cosmic [14] or Martincorena [15] and, on the other hand, for olfactory factors, which are believed to be mostly negative for cancer genes [35]. The results demonstrate that the *beta-binomial* and the *geometric* distributions dominates the best fit (Supplementary Figure 2A). If only the *betabinomial* and the *geometric* distributions were compared, 63% of the genes were best fitted using the *beta-binomial* (Supplementary Figure 2B). Moreover, for those genes best fitted with the *geometric* distribution, 98% would best fit the *beta-binomial* if the *geometric* were not considered whereas the genes best fitted with *beta-binomial* would not prefer the *geometric* (Supplementary Figure 2C). These results suggest that, overall, the best distribution tested is the *beta-binomial*.

### Hotspot detection algorithm

As shown above, the *beta-binomial* distribution seems to be a good model for most of the genes and it has been used to estimate recurrent alterations [36–38]. The use of a distribution is interesting because it provides the probability of observing *k* mutations allowing the possibility of assigning a p-value to biased amino acid positions (putative hotspots). Although a distribution could be a good model, the presence of hotspot mutations or biased sites would artificially increase the mutations counts at specific positions generating longer tails. This will generate deviations in the parameter values modifying the corresponding p-values and therefore falsely calling or not calling hotspots at uncertain conditions. To handle this, a mixed model is proposed having two components as *M* = *BetaBib*((*α,β*) +*F* where *M_k_* is the count of amino acid positions mutated *k* times. Without hotspots or deviated sites, the *F* vector is zero (all *F_k_*=0) and the number of amino acid sites mutated are explained entirely by the *betabinomial* component. This would generate very low differences between the observed and fitted distribution, which is measured by the Kullback-Leiber (KL) divergency (or *G-test*, see Methods). In the presence of hotspots or sequence biases, the KL divergency will be higher. Nevertheless, within the model, *F* can absorb the excess of amino acid positions at *k* mutations (*F_k_* > 0), providing a better fit for the *beta-binomial* and lowering the KL divergency. Therefore, the problem is to finding the optimal *a*, *b*, and *F*. For this, the devised step-wise algorithm, schematized in Figure 1B, first sets *F_k_*=0, then finds the most deviated amino acid positions at *k* mutations looking for lower values of cell scores. This is achieved exploring the possible combinations of *k* mutations and fractions of amino acid positions. In the example shown in Figure 1B, the first iteration finds *F_7_* = 1 while the second iteration finds *F_6_* = 1. The process ends because there is no sufficient improvement at the third iteration. In this way, the fitted *betabinomial*, conditioned to the fitted *F*, is more representative of most sites and mutations providing an unbiased estimation of the probability of *k* mutations at updated parameters *a* and *b*, which can be very different that without using the fixed effect at the start of the algorithm.

### Assessing the performance of the proposed algorithm

To objectively evaluate the performance of the proposed algorithm, simulations were used. The first simulation was performed assuming no hotspots. To simulate realistic scenarios, all genes were first fit to the *beta-binomial* without a fixed effect. Then, the observed *α_g_*, and *β_g_* values for 2,000 random genes *g* were used to generate positions distributions at the same number of the observed mutations. Finally, the proposed algorithm was run with this artificial data. The results show that the proposed algorithm has a specificity of 84.3% recognizing 0 hotspots when there are none (Figure 2A).

**Figure 2.**
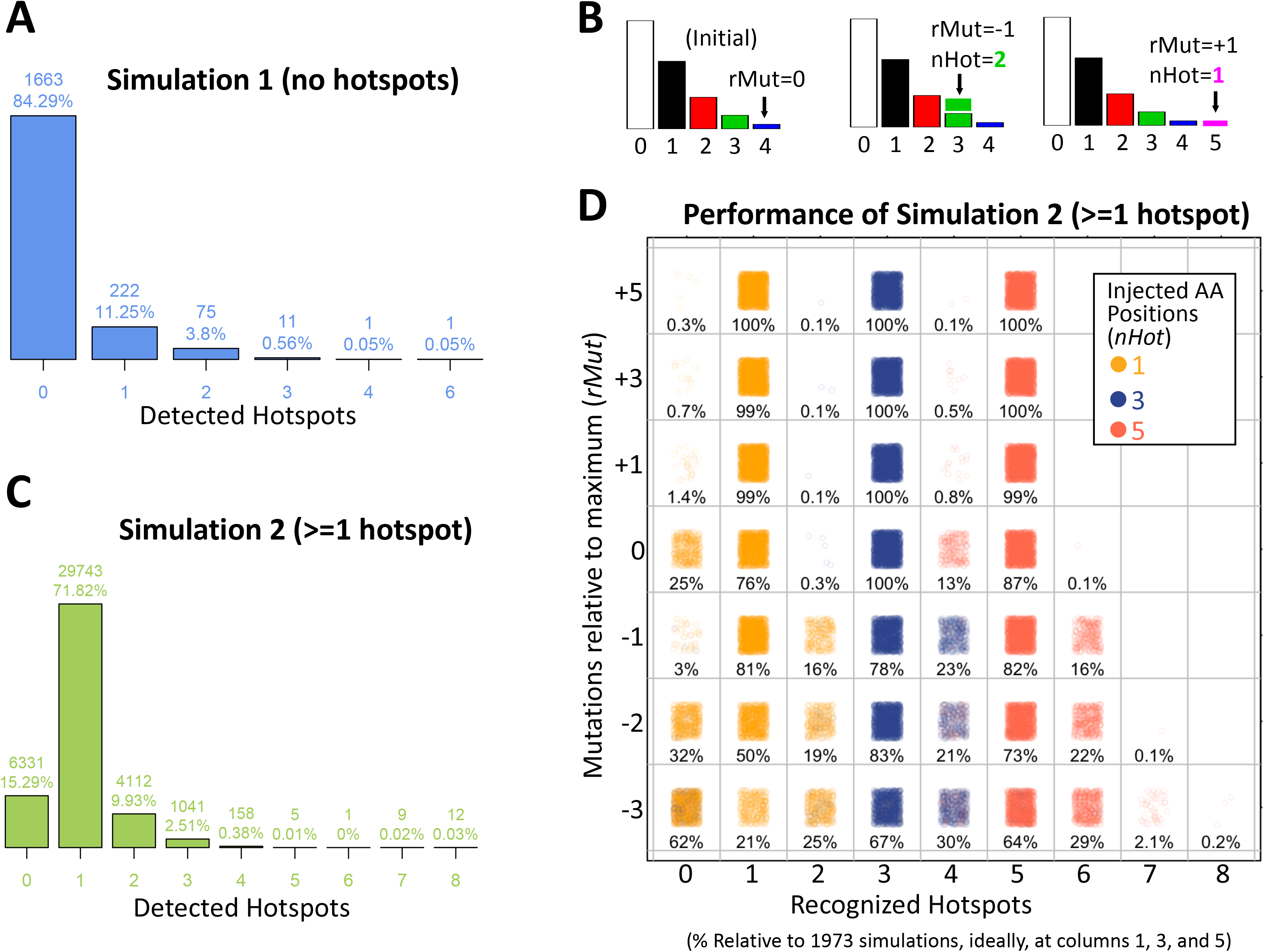
Performance of the proposed algorithm on simulated data. Panel A shows the distribution of the number of detected hotspots in simulation 1, which does not contain hotspots. Panel B shows how hotspots were injected. The left histogram shows original data. The middle and right histogram show the result of adding hotspots having 3 or 5 mutations respectively. The number of mutations is relative to the maximum number of observed mutations. In the example at left, the maximum is 4. Thus, *rMut=-1* refers to 1 mutation lower than 4 while *rMut=+1* refers to 1 mutation higher than 4. Finally, the *nPos* is the number of amino acid positions added to the specified number of mutations. Panel C shows the overall results of all simulations having hotspots. Here ‘detected hotspot’ stand for the sum of values > 0 from fitted F vectors. Panel D shows the performance of the algorithm depending on *rMut*. Each combination shows the percentage of simulated genes that showed the corresponding hotspots at the relative number of mutations.

The second simulation was performed assuming one or more hotspots (or biased amino acid positions). Note that the number of amino acid positions or the number of mutations is important because it could deviate far from the overall distribution or can be masked within dense regions of the distribution. For example, in Figure 1A, there is one hotspot carrying 6 mutations and another carrying 7 mutations, which are at +3 and +4 mutations farther than the last mutated ‘random’ mutation at 4. Similarly, in Figure 2B, two examples are shown. First, two hotspots are added having 3 mutations (relative to the maximum 4, these are at *rMut=-1*). Then one hotspot is added at 5 mutations (*rMut=+1)*. To generalize for any gene, for the simulations, the number of amino acid positions injected were *nHot*={1, 3, 5} whereas the number of mutations tested was *rMut*={−3, −2, −1, 0, 1, 3, 5} relative to the maximum number of observed mutations. In this way, injected hotspots at *rMut* <= 0 are harder to detect because are mixed with the overall distribution. Contrary, high values of *rMut* or larger *nHot* are easier to detect because the alteration has a deeper impact on the distribution. For these simulations, the same 2,000 genes employed in the first simulations were used. The results show that the proposed algorithm only fails to detect at least one hotspot in 15% of the simulations (Figure 2C). Thus, the algorithm has an overall sensitivity of 85%. Nevertheless, in more than 10% of the simulations, more than one hotspot was detected. To study the conditions of this behavior deeply, the performance of the algorithm for different *rMut* values was analyzed as shown in Figure 2D. The ideal well-known hotspots should contain more than the maximum random mutations, which corresponds to *rMut* > 0. The performance in these ideal hotspots was >= 99% for 1, 3, and 5 injected hotspots. If the hotspots are precisely the ones at the maximum number of mutations (*rMut=0*), the performance is 76% if there is only one hotspot, or close to 100% if there are 3 or more. If a hotspot is present but in the observed data is still below the maximum number of mutations (corresponding to *rMut < 0*), the performance decreases with both *nHot*, and *rMut* (Figure 2D). This scenario seems counterintuitive but because data in cancer has not been uniformly nor comprehensive acquired in all cancer types, it may be still useful if detected. In these cases, if only one hotspot is present, the overall performance decreases to 81%, 50%, or 21% corresponding to −1, −2, and −3 relative mutations and more than 15% of the times another false ‘hotspot’ is detected (see rows at *rMut= −1, −2,−3* and columns 2, 4, and 6). When three or five hotspots are present below the maximum mutations, the performance is in general higher but also increases the number of false ‘hotspots’ detected.

In summary, the proposed algorithm has an ideal performance (> 99% sensitivity and specificity) when the hotspots are those at the maximum number of mutations and the performance decreases with the number of hotspots or the relative position to the maximum number of mutations.

### Detecting hotspots in cancer data

From the proposed algorithm, the fixed effects *F* absorbs those positions that cannot be explained by the *beta-binomial* model alone. Thus, the fixed effect vector *F* mark hotspots while the fitted *beta-binomial* is able to, less biasedly, estimate its probability. The p-value was then corrected by a false discovery rate (FDR) approach [39]. Because potential hotspots are only those with a sensible number of recurrent positions, the FDR correction was estimated for sites whose recurrence were 4 or more. Only positions having FDR <= 0.01 were considered as hotspots. This was applied to TCGA mutational data, which includes 3,175,929 mutations from 10,182 patients across 33 cancer types (Supplementary Table 1). As a correction, hotspots were also called if the number of mutations were 9 or greater which includes many amino acid positions in *TP53*, *PIK3CA*, and *PTEN*, which result presumably to the overwhelming number of hotspots in these genes (Supplementary Figure 3). The detected hotspots are part of a database, *Hotspots Annotations*, available online (http://bioinformatica.mty.itesm.mx/HotSpotsAnnotations). Some representative examples of the hotspot detection are shown in Figure 3. For a well-known cancer gene, *EGFR*, 4 hotspots are clearly recognized carrying from 11 to 27 mutations. In addition, there were 4 AA positions carrying 5 mutations, 1 of 6 mutations, and 2 of 7 mutations that were effectively recognized by the algorithm but that were not significant under the above criteria after FDR correction.

**Figure 3.**
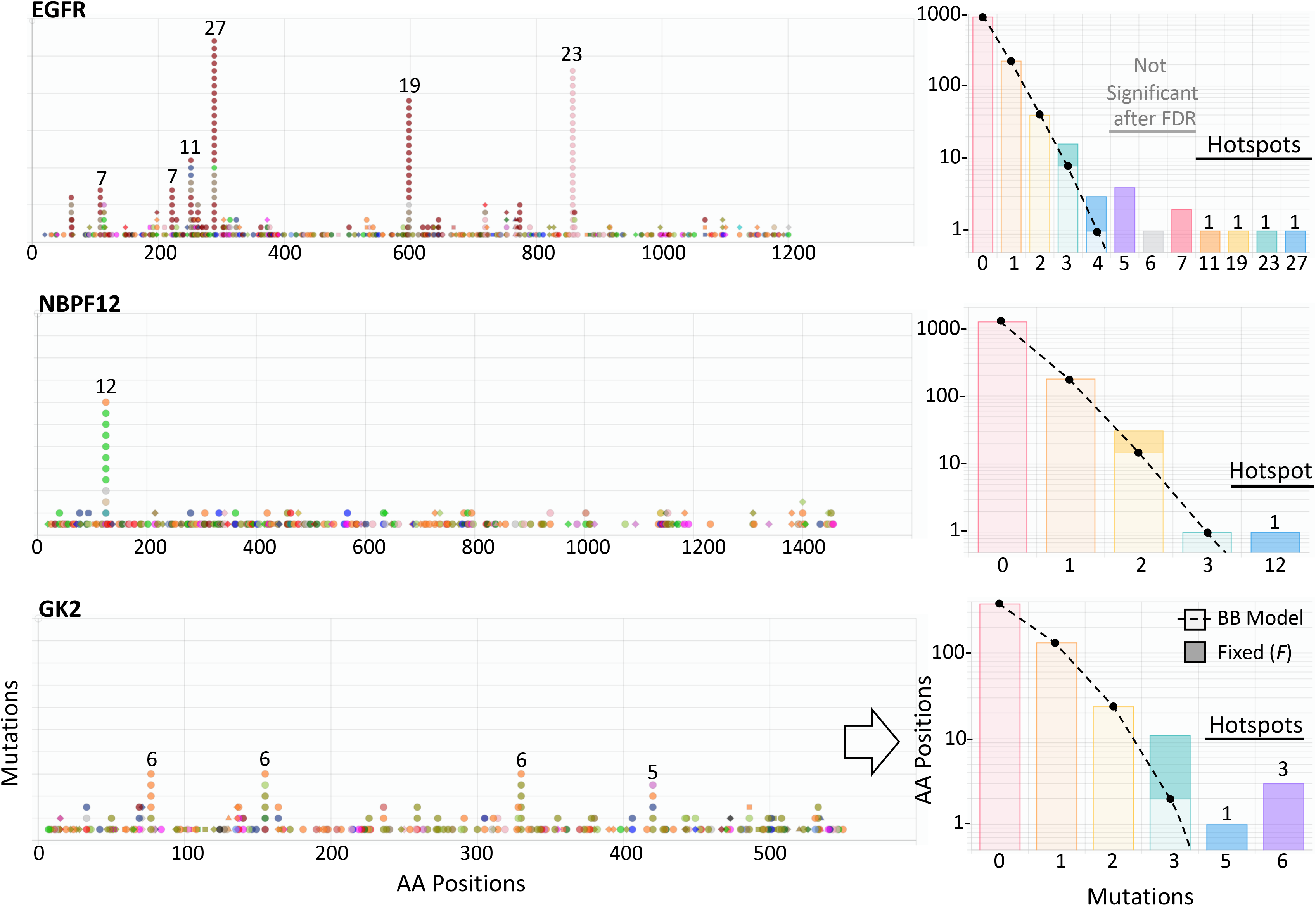
Examples of hotspots detections. Three examples of hotspots detections from TCGA data. The left drawings show the mutations along the protein sequence of three genes. Colors correspond to different cancer types. Symbols correspond to different types of mutations. The right histograms show the corresponding amino acid positions (vertical, in logarithmic scale) per number of mutations (horizontal). The beta-binomial component is represented in light bar colors and dotted line. The fixed effect is represented by darker bar colors. Significant hotpots are marked. Non-Significant fixed effects are also shown. Figures taken from http://bioinformatica.mty.itesm.mx/HotSpotsAnnotations developed in our research group.

Similarly, for *NBPF12* and *GK2*, not recognized as cancer genes in COSMIC, there were 1 hotspot accumulating 12 mutations in *NBPF12*, and 4 hotspots showing 5 to 6 mutations in *GK2*. In total, 3,860 hotspots were detected in 3,115 genes where 2,639 genes had only 1 hotspot, 378 genes contain 2 hotspots, and 98 genes showed 3 or more hotspots (Figure 4A). These hotspots cover 39,815 mutations representing 1.25% of the total mutations and 0.19% of the mutated sites. Common cancer genes showed many hotspots such as *TP53, PIK3CA, APC, PTEN, CDKN2A, ARID1A, FBXW7, NFE2L2*, and 6 or more were estimated in *ERBB2, CTNNB1, BRAF, CIC, KMT2D,* and *DNAH5*. *TTN* showed 7 ‘hotspots’ but has been marked repeatedly as a ‘false positive’ gene due to its size. The Table 1 shows the 96 genes showing 3 or more hotspots. This list is highly enriched in cancer genes, it contains 38% (n=37, p < 10^-53^) and 39% (n=38, p < 10^-31^) cancer genes from Cosmic [14] and Martincorena [15] respectively. Hotspots containing many mutations are commonly well-known because they have been spotted time ago such as IDH1 in gliomas, BRAF in thyroid, melanoma, and other cancer types. Nevertheless, an analysis of the distribution of mutations show high density corresponding to mutations between 5 and 9 reaching ~70% of detected hotspots (Figure 4B). This suggest that many hotspots are needed to be analyzed and experimentally studied.

**Figure 4.**
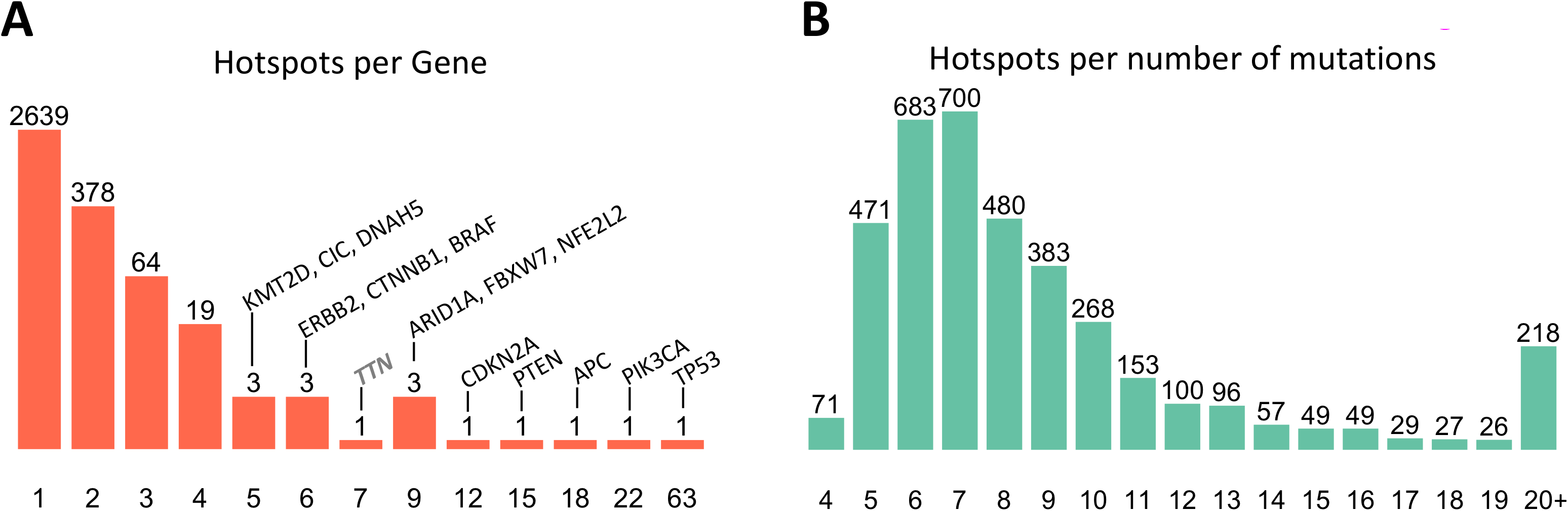
Distribution of hotspots per gene and mutations.

**Table 1.** Genes showing 3 or more recognized hotspots.

| Gene | HotSpots | Mutations<br>Min, Max | Gene | HotSpots | Mutation<br>Min, Max |
| --- | --- | --- | --- | --- | --- |
| BRAF | 6 | 11, 594 | KIAA2026 | 3 | 6, 11 |
| KRAS | 4 | 24, 564 | MDN1 | 3 | 7, 11 |
| PIK3CA | 22 | 9, 290 | PBRM1 | 3 | 9, 11 |
| TP53 | 63 | 16, 251 | PPM1D | 3 | 6, 11 |
| NRAS | 3 | 15, 203 | ZNF442 | 3 | 8, 11 |
| PTEN | 15 | 10, 112 | DNAH5 | 5 | 9, 10 |
| FBXW7 | 9 | 9, 69 | ATRX | 3 | 7, 10 |
| JAK1 | 3 | 14, 60 | C5orf42 | 3 | 7, 10 |
| CTNNB1 | 6 | 30, 50 | CSMD3 | 3 | 9, 10 |
| HRAS | 4 | 7, 50 | MECOM | 3 | 9, 10 |
| CDKN2A | 12 | 10, 41 | UVRAG | 3 | 5, 10 |
| APC | 18 | 9, 40 | TTN | 7 | 8, 9 |
| ERBB2 | 6 | 9, 40 | MGA | 4 | 7, 9 |
| PPP2R1A | 3 | 13, 33 | OR51S1 | 4 | 8, 9 |
| NFE2L2 | 9 | 7, 32 | ZNF292 | 4 | 6, 9 |
| ARID1A | 9 | 9, 29 | ADAMTS3 | 3 | 7, 9 |
| EGFR | 4 | 11, 27 | ADNP | 3 | 6, 9 |
| KMT2D | 5 | 9, 26 | ALB | 3 | 7, 9 |
| FGFR2 | 3 | 8, 25 | CHD1 | 3 | 7, 9 |
| SCAF4 | 3 | 10, 24 | DNAH7 | 3 | 9, 9 |
| SF3B1 | 4 | 7, 21 | RASA1 | 3 | 8, 9 |
| SPOP | 4 | 9, 19 | SAMD9 | 3 | 8, 9 |
| PIK3R1 | 4 | 10, 17 | TRIM23 | 3 | 6, 9 |
| KMT2B | 3 | 11, 17 | ZNF14 | 3 | 7, 9 |
| MBD6 | 3 | 12, 17 | ZNF732 | 3 | 7, 9 |
| MFRP | 3 | 6, 17 | KIF20B | 4 | 7, 8 |
| TPTE | 4 | 9, 16 | SLCO1B7 | 4 | 5, 8 |
| CD93 | 3 | 12, 16 | CWF19L2 | 3 | 7, 8 |
| KANSL1 | 4 | 8, 15 | FAM193A | 3 | 6, 8 |
| CHD4 | 3 | 11, 15 | OR2T2 | 3 | 7, 8 |
| CTCF | 3 | 11, 14 | UNC79 | 3 | 7, 8 |
| ZBTB7C | 3 | 9, 14 | ZNF502 | 3 | 6, 8 |
| CIC | 5 | 7, 13 | MSH6 | 4 | 6, 7 |
| NF1 | 4 | 8, 13 | VPS13C | 4 | 6, 7 |
| ARHGAP5 | 3 | 7, 13 | BTBD7 | 3 | 6, 7 |
| PRKDC | 3 | 8, 13 | CCDC27 | 3 | 7, 7 |
| SMAD2 | 3 | 7, 13 | CFAP61 | 3 | 7, 7 |
| YLPM1 | 3 | 7, 13 | GTF3C4 | 3 | 5, 7 |
| CASP8 | 4 | 9, 12 | PTPN11 | 3 | 5, 7 |
| ANK3 | 3 | 9, 12 | TDRD6 | 3 | 6, 7 |
| CNOT1 | 3 | 7, 12 | CCDC168 | 4 | 6, 6 |
| CNTNAP2 | 3 | 9, 12 | CSGALNACT1 | 3 | 5, 6 |
| MYOCD | 3 | 10, 12 | GK2 | 3 | 6, 6 |
| THSD7B | 3 | 10, 12 | PTPN13 | 3 | 5, 6 |
| ZFHX4 | 3 | 9, 12 | RALGAPA1 | 3 | 6, 6 |
| ALG13 | 3 | 8, 11 | RPS6KA5 | 3 | 5, 6 |
| C6 | 3 | 8, 11 | MAN2A1 | 4 | 5, 5 |
| DDX17 | 3 | 5, 11 | B3GAT2 | 3 | 5, 5 |
| HCN1 | 3 | 9, 11 | CLCA4 | 3 | 5, 5 |

### Variant types and sequence context in hotspots

Most hotspots methods focus on missense and nonsense mutations, which cover around 75% of all mutations. This has the advantage of focusing on clear biological effects but has the disadvantage of ignoring possible sequence biases that may help to recognize mechanistic effects. In addition, the proposed algorithm is inspired in estimating biases in the distribution of mutations along protein coding regions, which will be affected by selecting types of mutations. Therefore, all small mutations types were used. The disadvantage, however, is that not all variant types may show an interesting biological effect. In addition, it is known that hotspots may be focalized in specific sequence contexts [40]. Accordingly, a comparison of variant types and sequence contexts were performed between hotspots and the overall data in unique positions. To clearly expose the differences, only hotspots carrying 10 or more mutations were compared as shown in Figure 5 while the complete analysis is shown in Supplementary Figure 5. From the input data, the most frequent variant types are *missense, silent,* and *nonsense* accumulating 1.44, 0.564 and 0.116 million mutations. In hotspots, although the most frequent mutations are *missense* (n=750) surprisingly, *frame shift deletions* counts are very similar (n=742) even that *frame shift deletions* are more than 20 times less frequent in the overall data. *Frame shift insertions* were also high (n=327).

**Figure 5.**
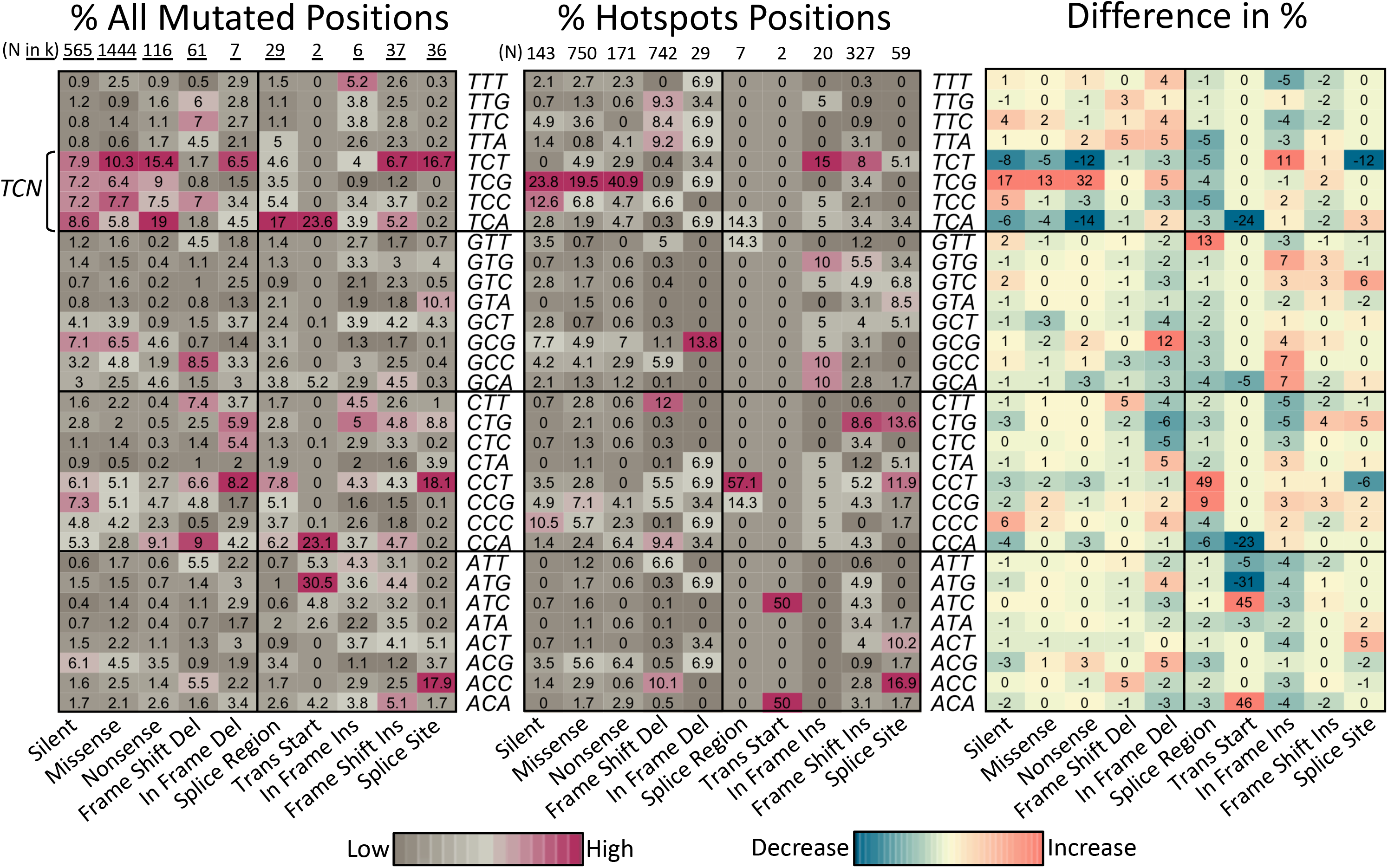
Comparison of mutated context sequences in hotspots. The left heatmap show the relative percentage of mutated positions per mutation type found in the whole dataset of TCGA data used. Only mutations types found in hotspots of 10 or more mutations are shown. Only distinct sites are considered. Total positions (N), are shown in thousands (k=1000). The heatmap at the middle shows equivalent percentages found at hotspots positions carrying 10 or more mutations. To facilitate interpretation, the heatmap at the right show the difference of the percentages. Fold-changes may vary. The Supplementary Figure 4 show details of other mutations types and for hotspots of 5 to 9 mutations.

The Figure 5 clearly show that while the sequence context T<u>C</u>N dominates the overall mutated positions mainly in the T<u>C</u>T sequence context (where the <u>C</u> marks the site of mutation), the T<u>C</u>G is by far the most recurrent context for hotspots while T<u>C</u>T, T<u>C</u>A, and T<u>C</u>C generally decrease. This pattern seems to be clearly present in *missense* and *nonsense* and partially also in *silence* mutations suggesting that there is some type of preference or selection for the T<u>C</u>G context in these types of variants. Similarly, for hotspots carrying 5 to 9 mutations, the T<u>C</u>G increase is also observed (Supplementary Figure 4). However, in these hotspots, an increase in G<u>C</u>G, then C<u>C</u>G and A<u>C</u>G, were also present suggesting that the overall preference for 5 to 9 mutations seems to be xN<u>C</u>G. All these results concur with the pattern of mutations from APOBEC [40]. For *frame shift deletions* the observed differences are not so strong, suggesting that, overall, selection pressure is absent or low. The highest increases in differences (+5 relative %) were in A<u>C</u>C, C<u>T</u>T, and T<u>T</u>A. For other types of variants, the changes or the number of occurrences in hotspots are low.

### Hotspots across cancer types

It is known that cancer types differ in the frequency of mutations per gene [35]. It has also been proposed that driver mutations may accumulate from 1 to 10 depending on the cancer type [15]. Therefore, a comparison of hotspots across cancer types were performed. First, it was noted that the percentage of samples not carrying any hotspot mutation formed three to four clusters of cancer types (Figure 6A), which also correlated with the overall mutation rate. The clusters include more than 60% of samples (TGCT, KIRP, KIRC, MESO, PCPG, PRAD, KICH, and ACC), then between 25% and 60% of samples (THCA, THYM, OV, GBM, BRCA, CESC, LAML, DLBC, LIHC, SARC, CHOL), those between 10% and 25% (PAAD, LGG, LUSC, HNSC, ESCA, LUAD, BLCA, STAD), and those below 10% (SKCM, COAD, UCEC). UVM, UCS, and READ show also low percentage of samples not carrying hotspots but its distribution is more similar to one of the first three clusters. STAD, SKCM, COAD, and UCEC show around 20% or more samples carrying 10 or more hotspots, which is also consistent with the high rate of mutations of these cancer types. It is well known that TP53, PIK3CA, and RAS gene family show recurrence in many cancer types but others genes are more specific. For example, IDH1/2 in gliomas, AKT1 and GATA3 in BRCA, SPOP in PRAD, and BRAF in THCA. Therefore, three approaches were performed to highlight cancer-specific hotspots. First, the top 10 most frequent hotspots per cancer type were estimated as shown in Table 2. Beside the above cancer-specific genes, other high frequent hotspot can be noted such as GTFI2 in THYM, GNAQ in UVM, CTNNB1 in LIHC, VHL in KIRC, CDKN2A in HNSC, and NFE2L2 in LUSC. Second, an analysis of the number of cancer types per hotspot shows that most hotspots (91%) are formed by mutations from 2 to 6 cancer types (Figure 6B). Thus, only 95 hotspots (2.46%) are strictly cancer type-specific (Figure 6C). For example, VHL p.158 in KIRC, APC p.935 in COAD, and CDH1 p.23 in BRCA. Third, because of these results, for each hotspot the major cancer type was calculated. Then, if its contribution to the total number of mutations were higher than 50% or if it were higher than 25% and the number of mutations were higher than 10, it was selected as ‘cancer-enriched’. Thus, the number of hotspots per cancer type was very high for UCEC, STAD, SKCM, and COAD as shown in Figure 6C, presumably due to high mutations rates. The Table 3 shows the hotpots for the rest of cancer types and the complete list is shown in Supplementary Table 1. This is interesting because it highlights genes not well studied such as NBPF12 in BRCA, LPAR6 or ASXL2 in BLCA, and FGGY in LUSC, which is being studied recently [41].

**Figure 6.**
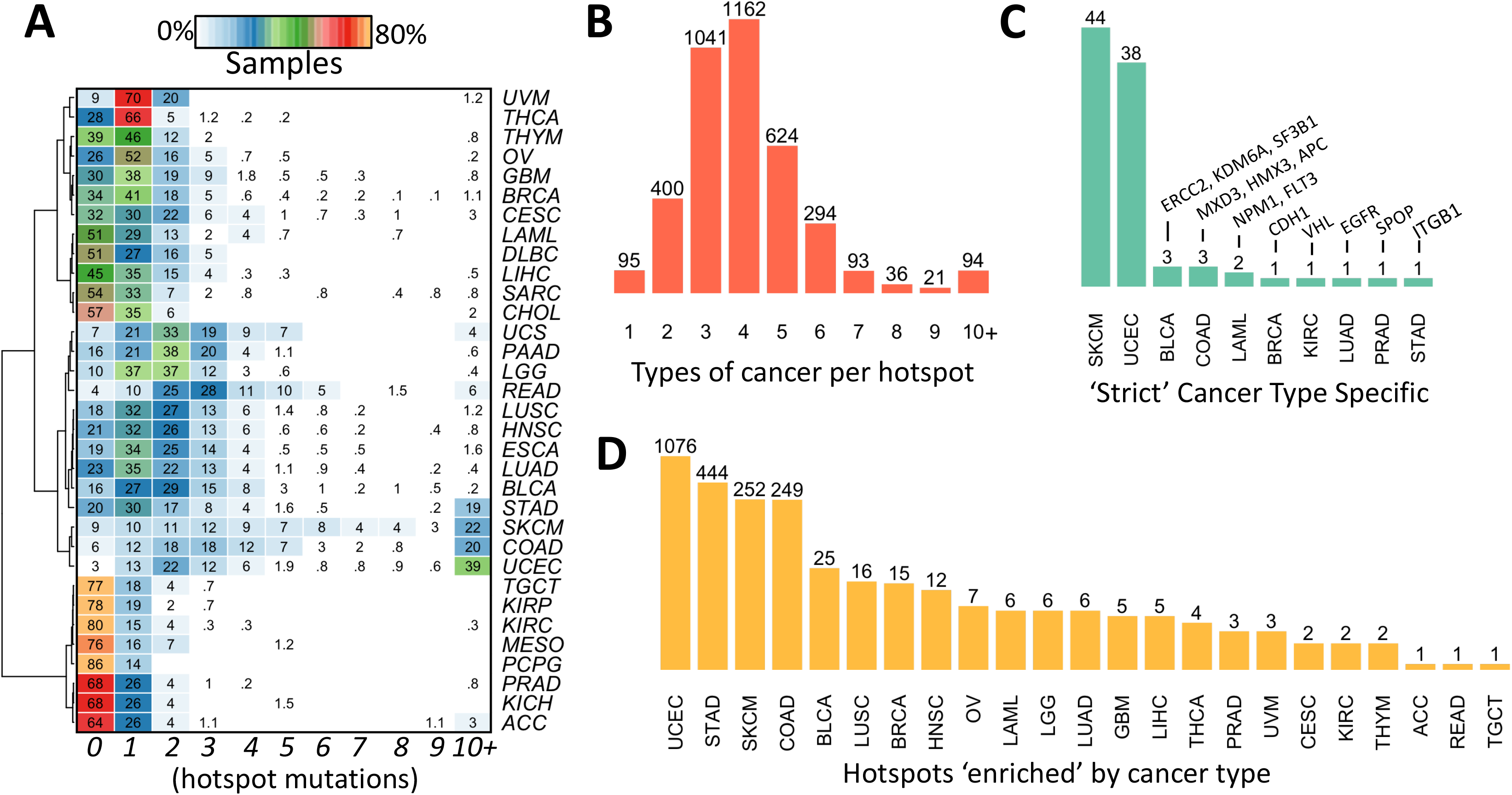
Distribution of hotspots across cancer types. Panel A shows the percentage of samples containing *k* hotspots. Panel B shows the different types of cancer per hotspot. Panel C shows the 95 hotspots found at only one cancer type. Supplementary Table 2 shows the genes that are strict cancer type-specific. Panel D shows the hotspots that are majorly represented by one cancer type. Supplementary Table 3 shows the genes that are enriched by cancer type.

**Table 2.**
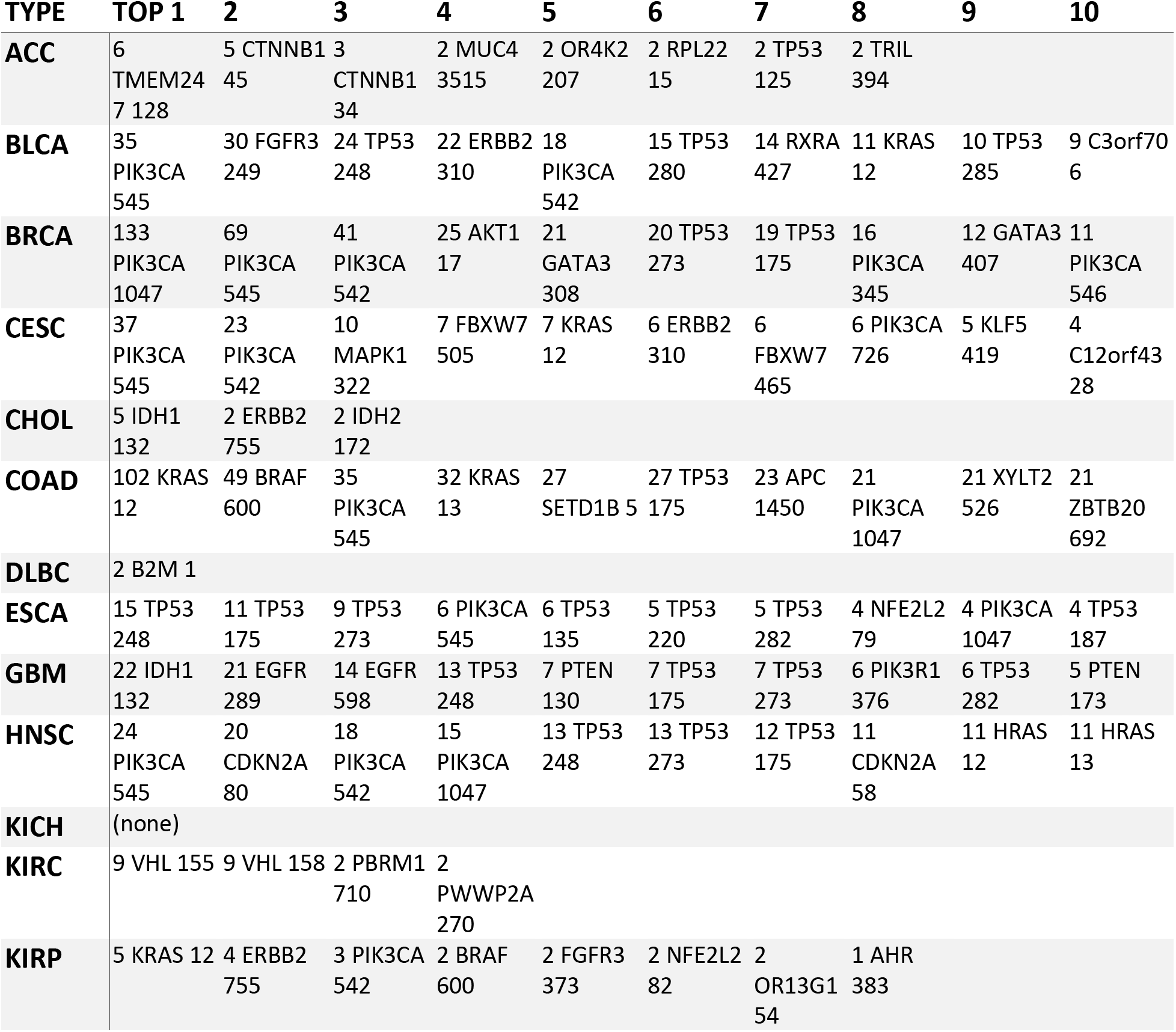
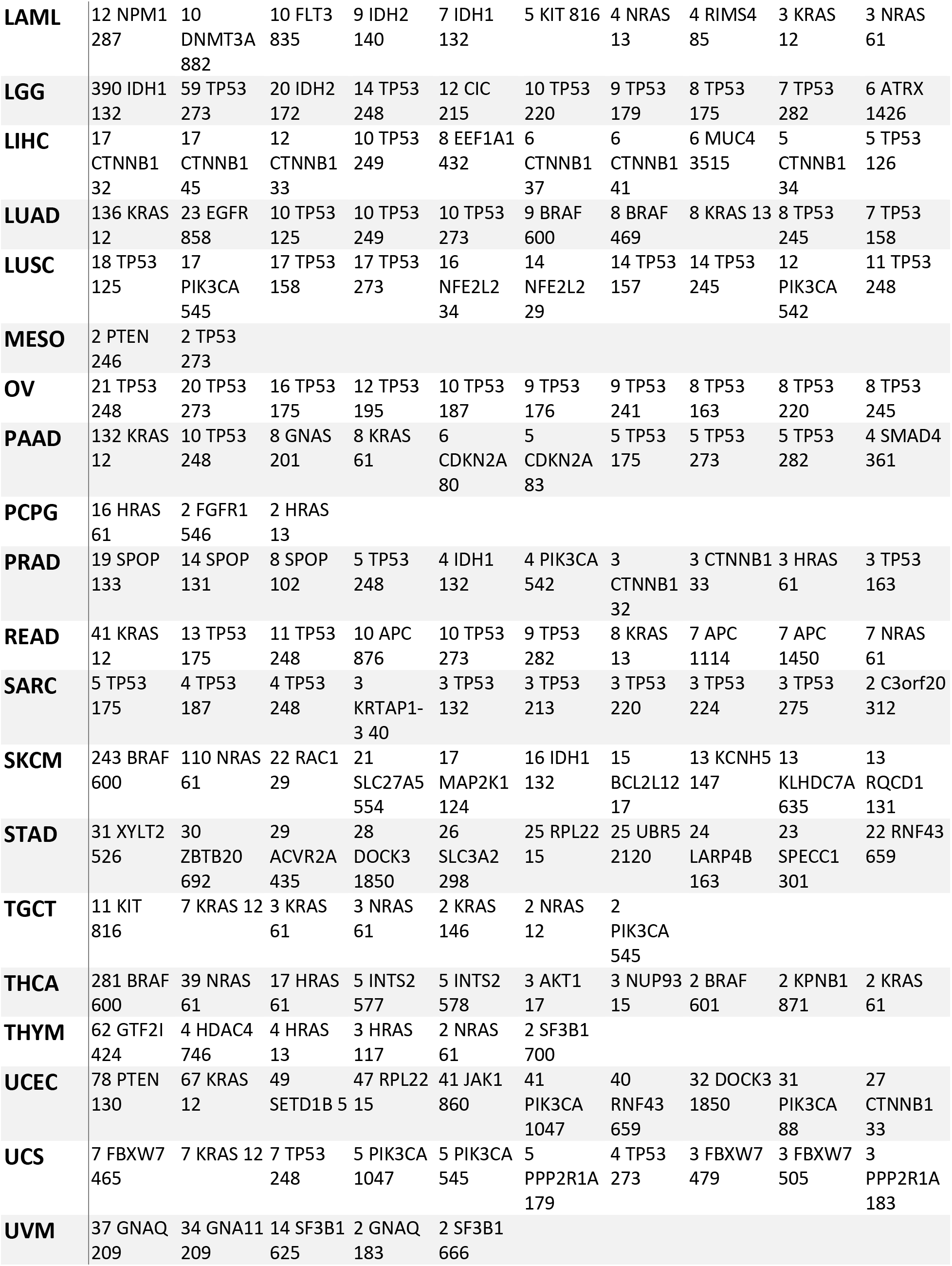
Top 10 most frequent hotspots per cancer type (# patients GENE position).

**Table 3.**
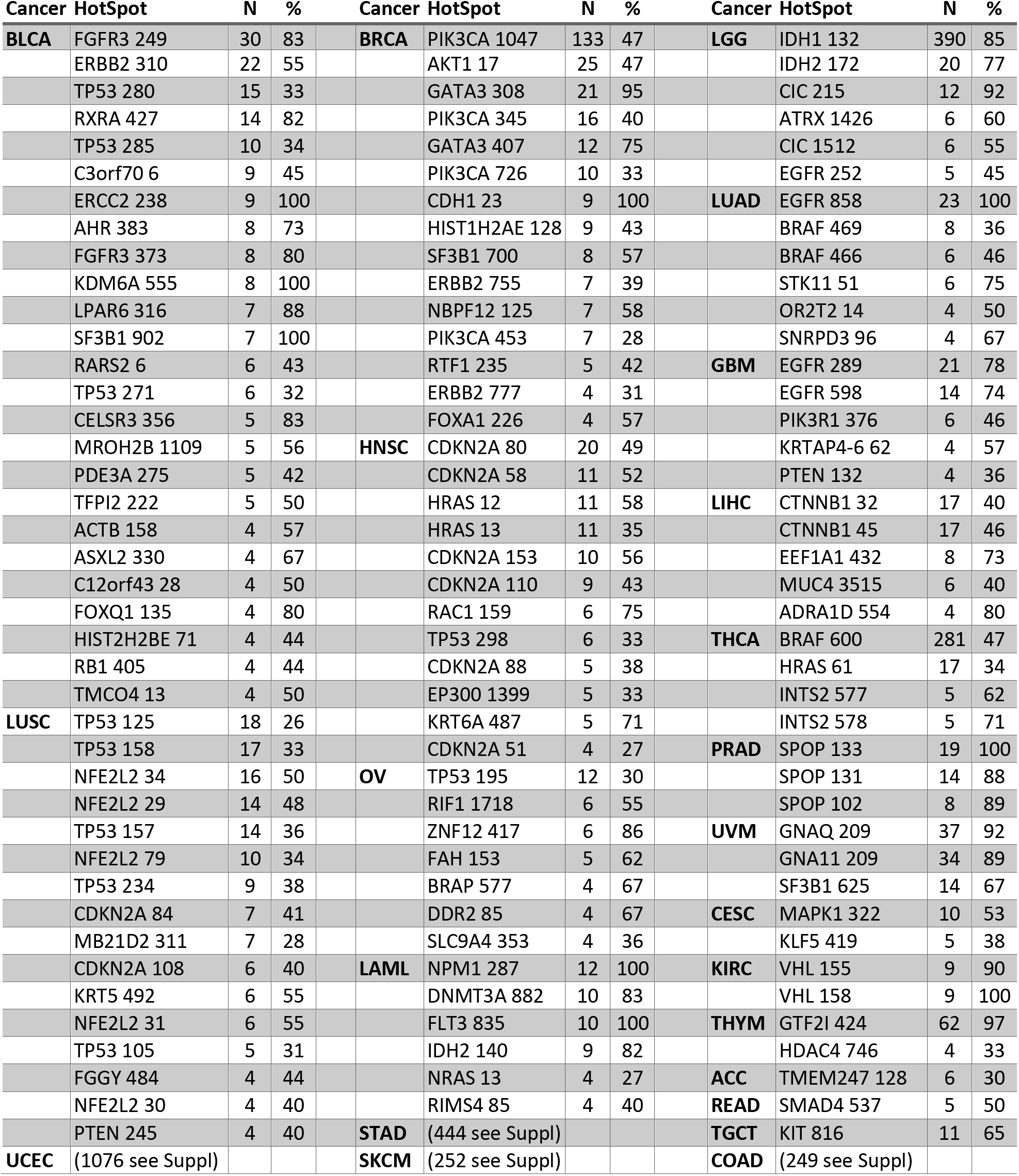
Cancer enriched hotspots.

### Model parameters correlates with background mutation rates

The estimation of background mutation rates is important for mutation detection methods because it helps to determine deviations [42]. Instead of the expected number of mutations, the fitted *beta-binomial* model can be used to provide estimations of the probability of *k* mutations along chromosomes. By definition, contiguous genes should show similar probabilities even that the fitting was independent. Small deviations of an overall probability should highlight important genes and systematic deviations should show artifactual genes or regions. To validate this, the estimated p-values were compared between genes along chromosomes. The Figure 7 shows a representative example of the estimations for the chromosome 1 (Supplementary Figure 4 shows all chromosomes) for the p-value of 0 and 1 mutations (shown in black and red respectively). It is clear that the smoothed mean show some peaks that colocalize with olfactory receptors (vertical gray lines), which has been shown to be highly correlated to late replication timing, low expression, and higher mutation rates [35]. Other gene clusters can be identified, for example, late cornified envelope (LCE) gene cluster in Chr1 (Figure 7), regenerating family member (REG) in Chr2, protocadherin beta gene cluster (PCDHB) in Chr5, the histone 1 cluster in Chr6, among others (Supplementary Figure 5). Specific deviations such as CDKN2A in Chr9, PTEN in Chr10, TP53 in Chr17 among other are also visible (Supplementary Figure 5). These results show that the proposed algorithm provides consistent estimations.

**Figure 7.**
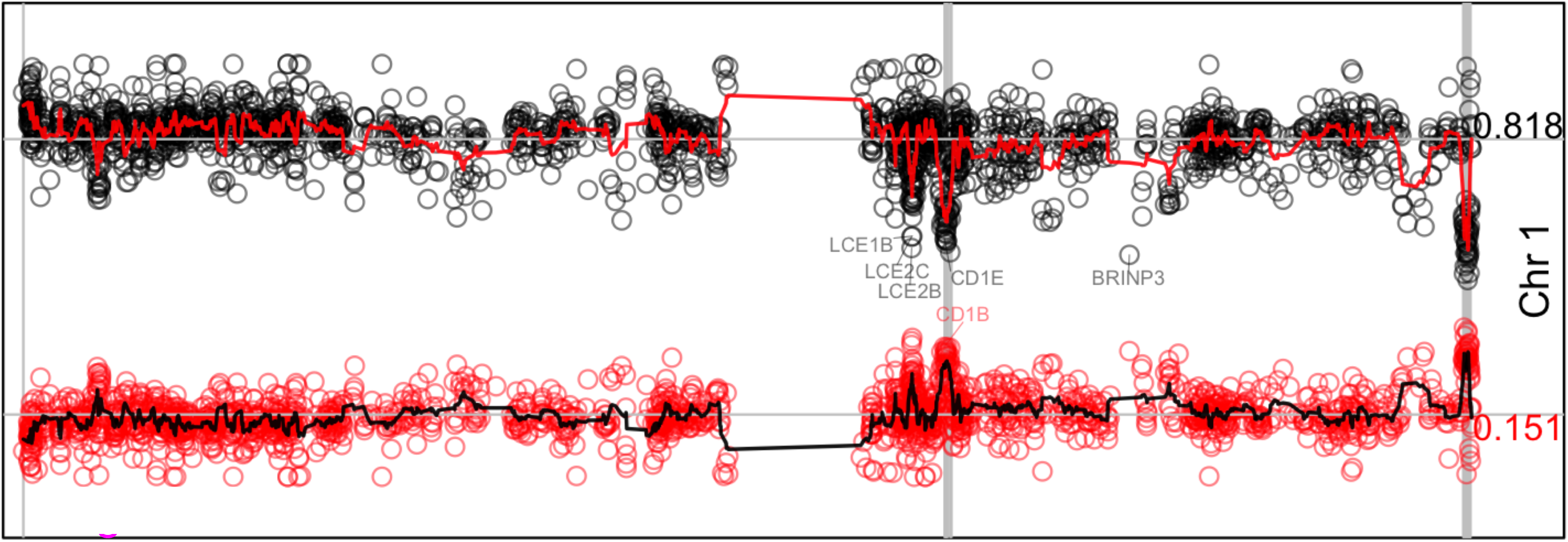
Model estimations along chromosome 1. The figure shows the density estimations of 0 mutations (dots in black) and 1 mutation (dots in red). The red line in top and black line in bottom show the smoothed estimation (window=5). The mean value, 0.818 for the former and 0.151 for the last, is shown at right and represented by a horizontal gray line. Vertical gray lines represent genomic positions for annotated olfactory receptors. Some genes farther than 3 standard deviations are annotated. Supplementary Figure 5 shows equivalent information for all chromosomes.

## Discussion

This manuscript shows an algorithm to identify highly recurrent mutations at specific amino acid positions in cancer. The algorithm fits the distribution of amino acid positions along number of mutations using a mixed model that includes a beta-binomial model plus a fixed effect (Figure 1).

The comparisons of different distributions lead to select the *beta-binomial* model. This makes sense because, in principle, the mutation can be seen as a binomial process during replication and/or repair. Then, instead of fixing *p* along the gene in the binomial process, *p* is random drawn from a *beta* distribution, which absorbs uncertainty due to patient, different positions, and sequence contexts resulting in allowing more uncertainty, covering observed overdispersion, and fitting the data better. Other statistical models could be tested but the justification, the interpretation, and the adequacy of the model may be difficult.

One of the problems when proposing a predicting or discovery algorithm is how assessing the accuracy. Although other algorithms and models have been proposed, most of them use lists of positive and/or negative curated genes as benchmarking. Instead, simulations were used here showing that overall the sensitivity and specificity was ~85%. More importantly, in conditions common for hotspots such as at highest number of mutations, the algorithm shows accuracies around 99%.

Few genes such as TP53, PIK3CA, and PTEN, showed a different tendency in fitting than most genes (Supplementary Figure 3). This is presumably due to the high number of hotspots and mutations backed up by the observation that closer genes such as CDKN2A, GATA3, and APC also show high hotspots. This is not a problem because these are well-known cancer genes. Nevertheless, it would be interesting to observe other genes once more mutation data is aggregated in the coming years.

It is assumed that a hotspot have functional impact in cancer [1]. Nevertheless, recent advances have shown that many hotspots arise by artifacts in local sequences such as hairpins susceptible for APOBEC enzymatic activity [40], including the detected gene MB21D2. Therefore, it is difficult to confirm in advance which hotspots will be functional. However, the first step is to detect those that under a certain model seems to be potential hotspots. These hotspots are provided here. Thus, how hotspots must be selected for functional validation? First, those that are well-known cancer genes whose hotspot have not been experimentally tested. Second, the genes showing many hotspots or high number of mutations at the hotspot. These would provide further certainty that any of its hotspots are indeed functional.

Nevertheless, in the analysis of cancer data, most genes only show 1 hotspot and most hotspots were found supported by less than 10 mutations (Figure 4). Third, check that the gene has not been listed for APOBEC activity [40]. In this context, the database HotSpotsAnnotations has been created (http://bioinformatica.mty.itesm.mx:8080/HotSpotsAnnotations) which has been annotated for APOBEC, the ratio of non-synonymous by synonymous mutations, and can be manually annotated by the research community [43]. Fourth, further verification is needed if the gene is super-sized or within artifactual regions such as those around olfactory receptors. Fifth, check the criteria of the ratio of non-synonymous to synonymous mutations [15]. Finally, frame shifts deletions and insertions have not been well studied in the hotspot context and in statistical models. Around one third of the detected hotspots included these mutation types.

The observation that *T<u>C</u>G* is more prone to form hotspots does not seem to be due to the lack of covariates in the model used. This is based on the fact that sequence context in hotspots were analyzed after normalization by percentage comparing the observed mutational spectra and the hotspots. That is, if all mutational contexts would have similar probability of being established as a hotspot, similar percentages would be observed in hotspots. Instead, more than two-fold was observed in *T<u>C</u>G* for single nucleotide variants.

Most hotspots carry between 5 to 9 mutations (70%) and also are formed by mutations of different cancer types (91%). Therefore, many hotspots were only detected when mutation from all cancer types were aggregated highlighting the importance of integrating databases. Consequently, as more mutation data is accumulated, more precise detections can be done. One issue is that all datasets must be processed in compatible pipelines, genome annotations, and transcripts to avoid inconsistencies. In this context, other databases such as those from the International Cancer Genome Consortium (ICGC) should improve and confirm the results.

## Conclusion

Simulations of the proposed algorithm that fit a mixed model of *beta-binomial* plus a fixed effect demonstrated excellent performance for hotspots at highest mutations (around 99% accuracy) and acceptable overall performance (85%). The algorithm was applied to TCGA cancer data detecting more than 3,860 hotspots after FDR correction that account for around 1.25% of the total number of mutations and 0.19% of the mutated amino acid sites.

## Acknowledgements

I thank Dr. Jose Tamez, Dr. Emmanuel Martinez, and all participants in the Bioinformatics seminar for their comments and recommendations.

## Supplementary Material

**Supplementary Figure 1.**
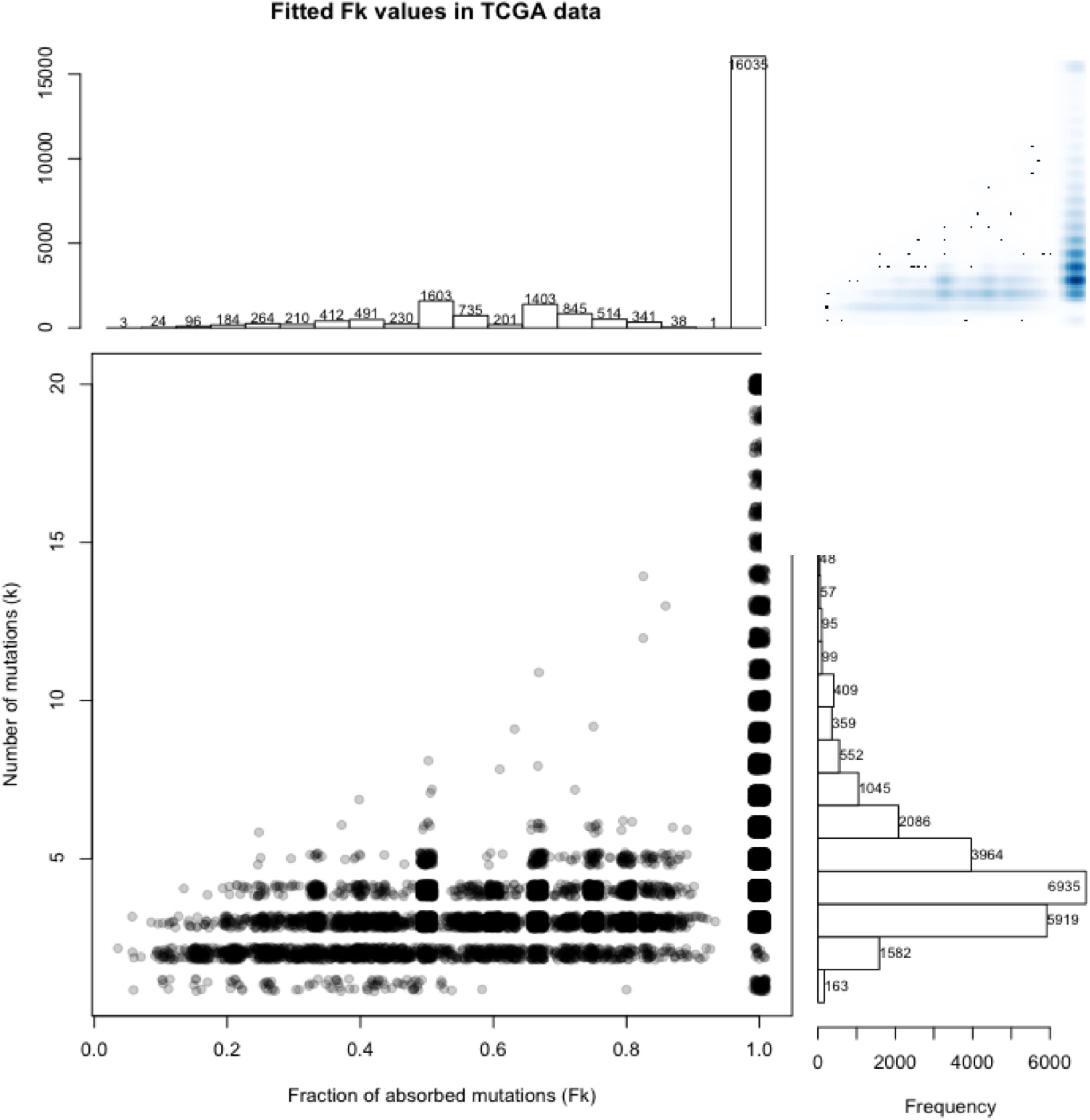
Fitting characteristics in TCGA data. The figure shows the distribution of the fitted fixed effect F (horizontal) relative to the total number of mutations (vertical). A value of 1 represent 100% of the mutations. It is clear that most fittings absorb 100% of mutations (highest vertical bar at top). Nevertheless, few fittings are fractions of the total number of mutations, mainly at low number of mutations (5 or lower). Therefore, these fittings capture biases in the distribution relative to the beta-binomial model.

**Supplementary Figure 2.**
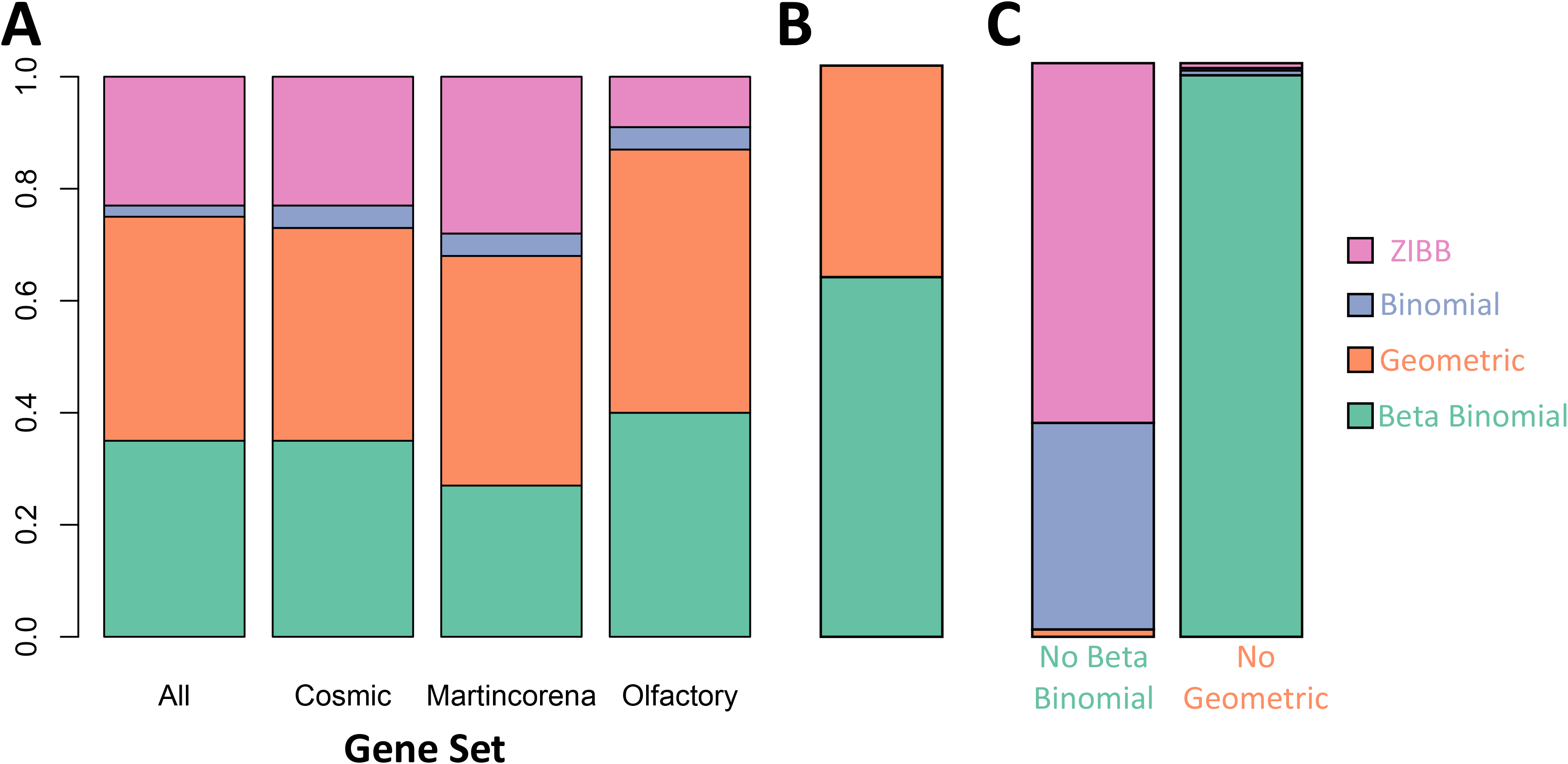
Comparisons of best fitting among four distributions. Panel A shows a comparison of the best distribution for different sets of genes. Panel B compares the best two distributions. Panel C shows results if one distribution is not considered.

**Supplementary Figure 3.**
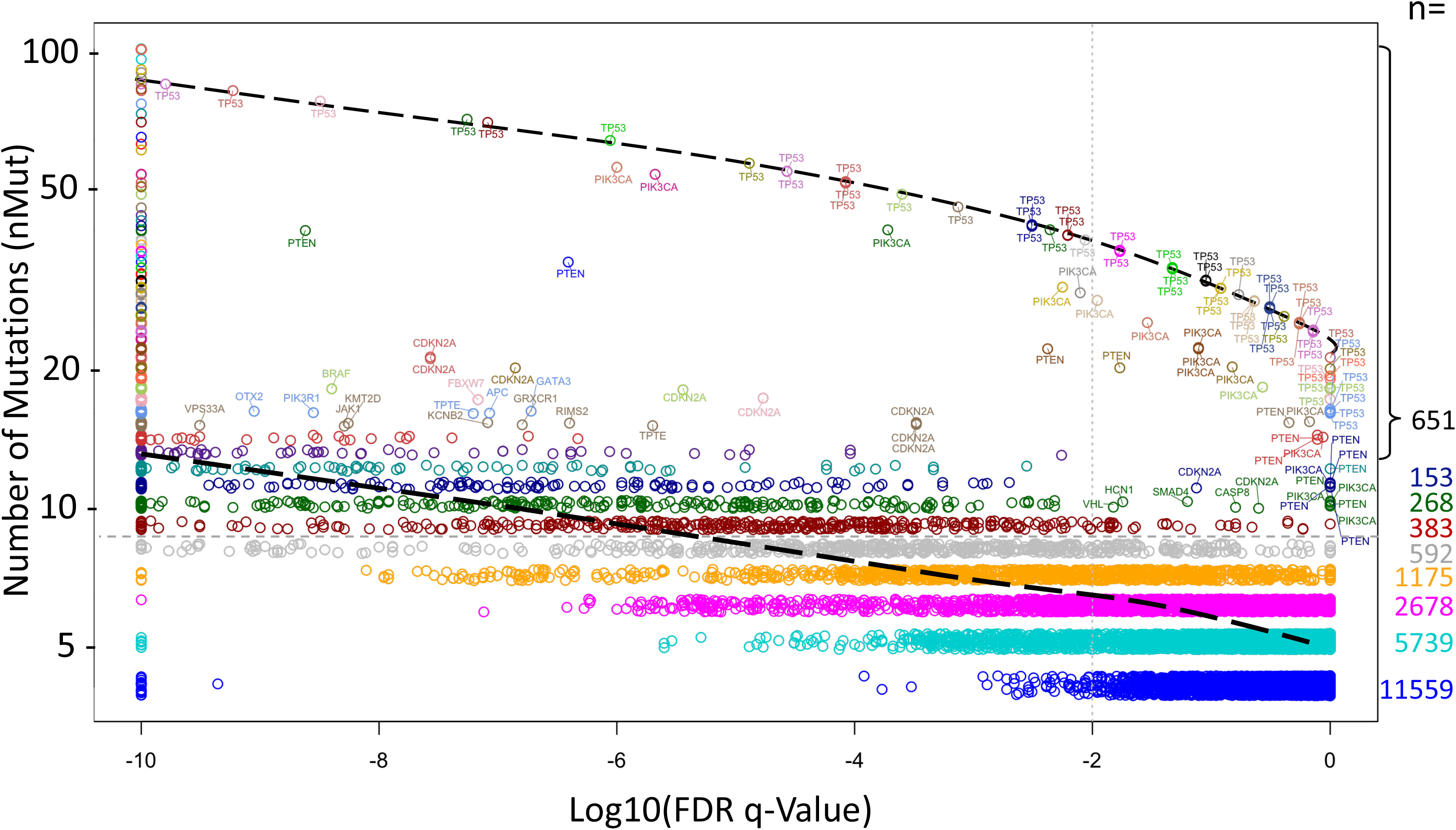
FDR estimation for amino acid positions carrying 4 or more mutations. Two tendencies are observed, a major below ~20 mutations and a minor above ~20. The last includes TP53, PIK3CA, and PTEN hotspots mainly as labelled. The vertical *nMut* was jittered for clarity in density estimation. FDR q-value was limited to 10^-10^ (at left) for clarity.

**Supplementary Figure 4.**
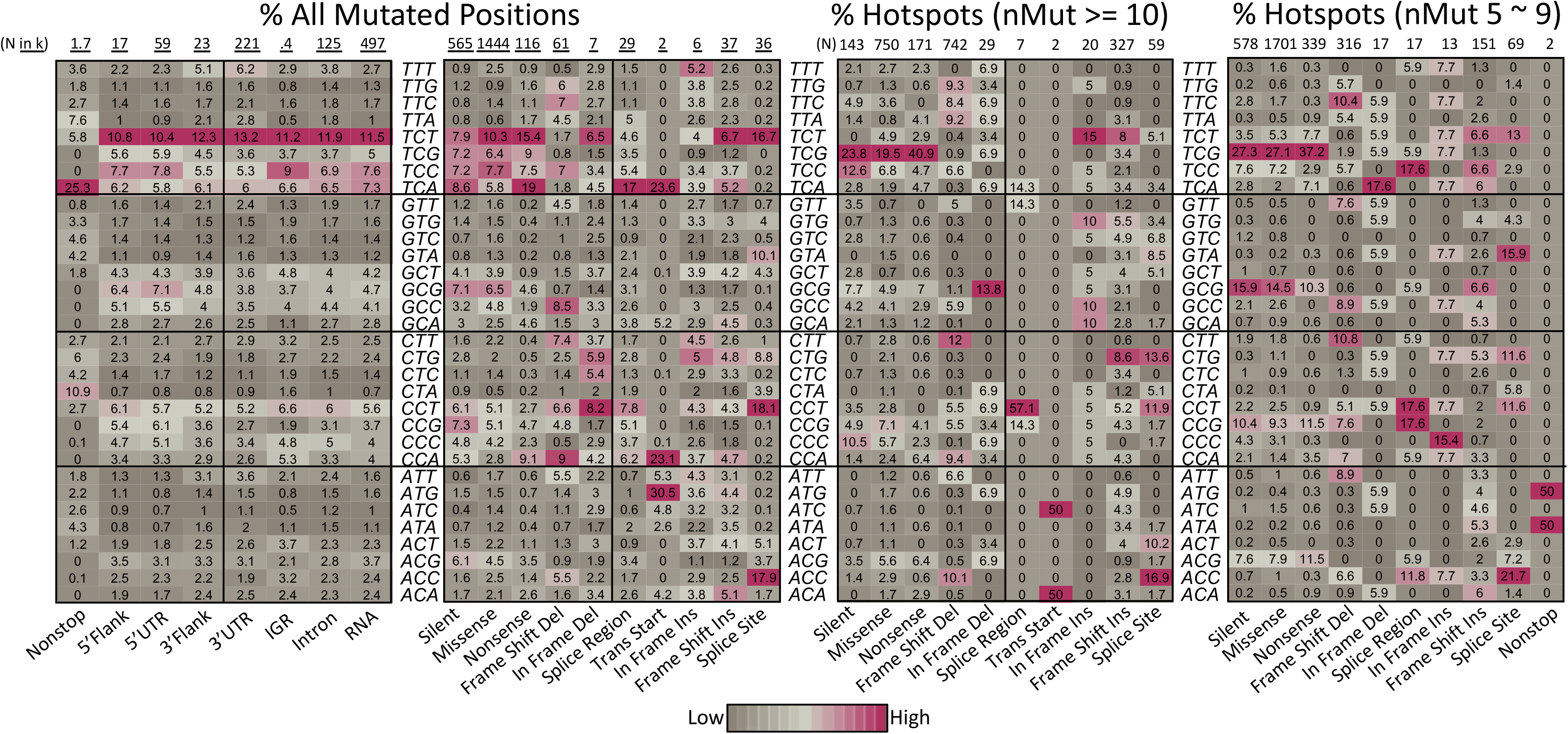
Comparison of mutated context sequences in hotspots. The first two heatmaps show the relative percentage of mutated positions per mutation type found in the whole dataset of TCGA data used. The first show the types of mutations not found in hotspots of 10 mutations or more. The second show the types of mutations found in hotspots of 10 or more mutations. Only distinct sites are considered. Total positions (N), are shown in thousands (k=1000). The third heatmap shows equivalent percentages found at hotspots positions carrying 10 or more mutations. The last heatmap at right show equivalent percentages for hotspots carrying 5 to 9 mutations.

**Supplementary Figure 5.**
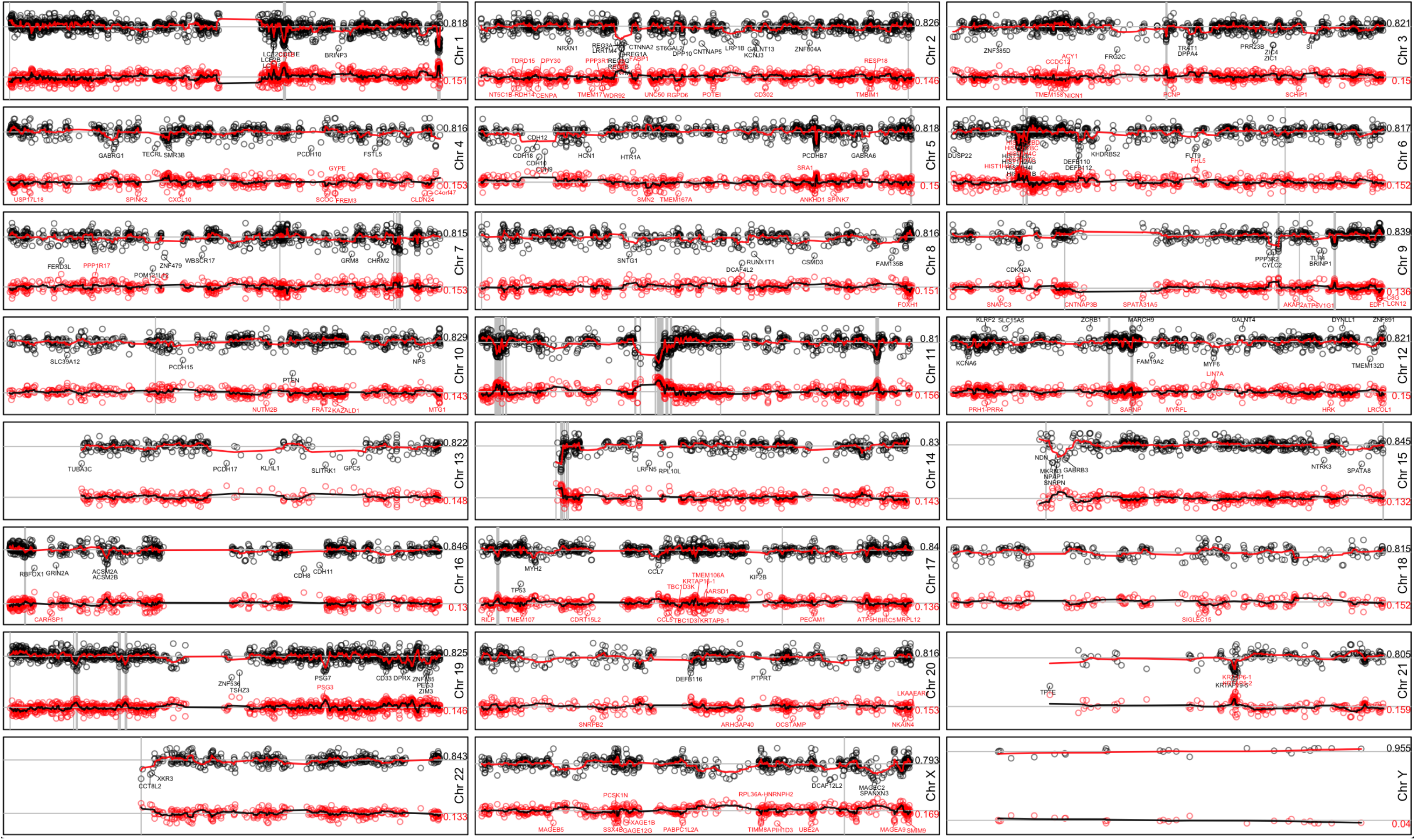
Model estimations along chromosomes. Each panel shows the density estimations of 0 mutations (dots in black) and 1 mutation (dots in red) for a chromosome (as labeled at right). The red line in top and black line in bottom show the smoothed estimation (window=5). The mean value is shown at right and represented by a horizontal gray line. Vertical gray lines represent genomic positions for annotated olfactory receptors. Some genes farther than 3 standard deviations are annotated.

**Supplementary Table 1. Number of samples mutations per cancer type used in this study.**

**Supplementary Table 2. Genes strictly cancer type-specific.**

**Supplementary Table 3. Genes enriched by cancer type-specific.**

